# Seizure Circuit Activity in the Theiler’s Murine Encephalomyelitis Virus Model of Infection-induced Epilepsy Using Transient Recombination in Active Populations

**DOI:** 10.1101/2025.05.20.655185

**Authors:** Alexandra N. Petrucci, Ted Abel, Karen S. Wilcox

## Abstract

Epilepsy affects one in twenty-six individuals. A major cause of epilepsy worldwide is viral encephalitis. Central nervous system infections can provoke seizures in the short term and increase the risk of spontaneous, recurrent seizures post-infection. However, the neural mechanisms underlying seizures during acute infection are unknown. These neuronal changes can be studied in C57BL6/J mice infected with Theiler’s murine encephalomyelitis virus (TMEV). TMEV-infected mice experience seizures 3-8 days post-injection (DPI), clear the virus by DPI 14, and may develop chronic, acquired temporal lobe epilepsy. TMEV may incite seizures during the acute infection period through inflammation, reactive gliosis, and cell death in hippocampal area CA1. Here, we explore the neuronal circuits underlying acute seizures in TMEV-injected mice using c-Fos driven TRAP (targeted recombination in active populations). TRAP mice (c-Fos- CreERT2 x CAG-tdTomato) were injected with PBS or TMEV and gently handled on DPI 5 to induce seizures. 4-OHT was administered to mice either 1.5 or 3 hr after seizures to tag the active cells expressing c-Fos with tdTomato. After 1 week, the mice were sacrificed and whole mouse brains were sectioned and immunostained for tdTomato expression. Percent area of fluorescence was quantified, and comparisons were made between TMEV-injected mice and PBS controls, sites ipsilateral vs contralateral to TMEV injection site, and between sexes. TdTomato expression was elevated in the TMEV-injected mice in the ipsilateral and contralateral hippocampus, thalamus, lateral septal nucleus, basal ganglia, triangular septal nucleus, fornix, and corpus callosum. Critically, the expression pattern suggests that seizures induced on DPI 5 arise from the hilus, dentate gyrus, and CA3 hippocampal subregions. Generalized seizures during acute TMEV infection may have propagated to the contralateral hemisphere via CA3 and the hippocampal commissure. TRAP has not been previously utilized in the TMEV mouse model and these experiments address crucial questions regarding seizure spread during TMEV infection.

**GRAPHICAL ABSTRACT:** 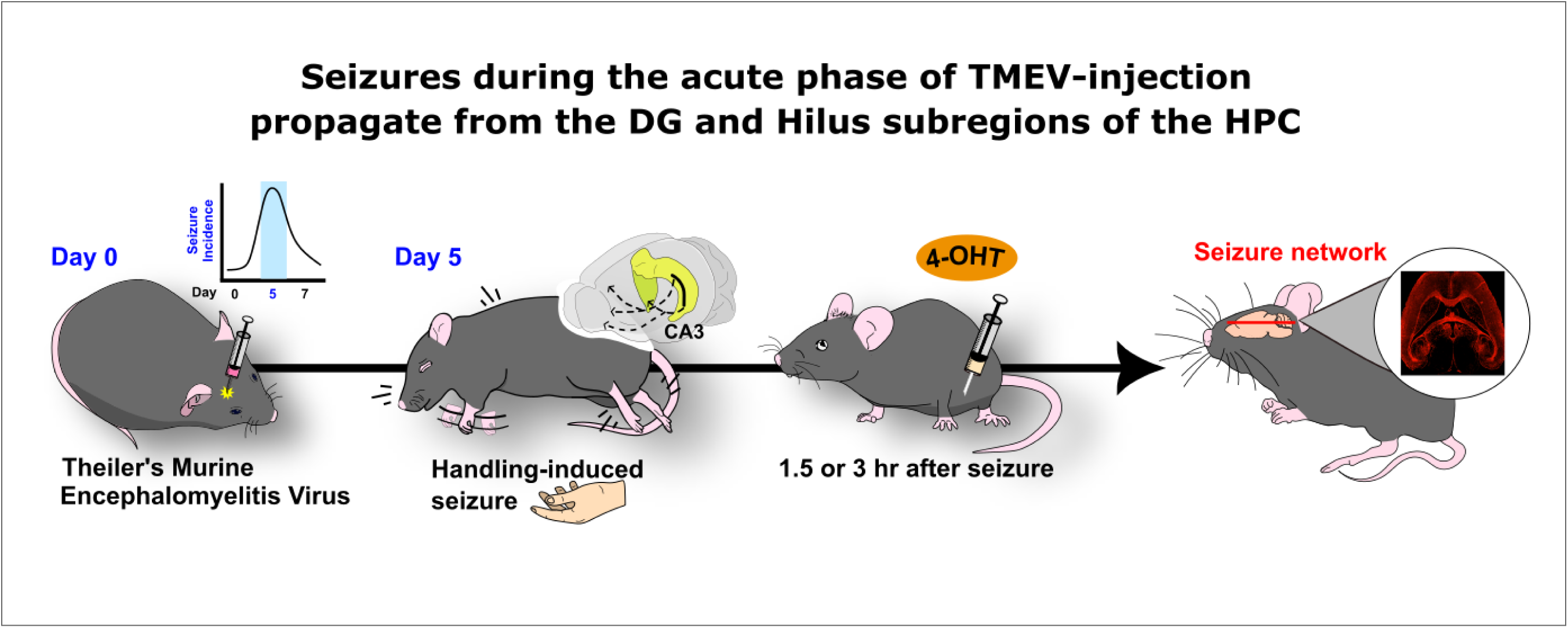

## INTRODUCTION

Viral-induced encephalitis is a major cause of epilepsy globally^1^. Over 100 viruses are known to cause epileptic encephalitis in humans^2^. Inflammation and viral-induced damage provoke seizures during active infection and greatly increase risk of developing spontaneous, recurrent seizures ^1,3^. Patients that experience seizures during acute infection are 22 times more likely to develop chronic epilepsy than the general population^4^.

One useful model of infection-induced temporal lobe epilepsy (TLE) develops when C57BL6/J mice are injected with the Daniel’s strain of Theiler’s murine encephalomyelitis (TMEV) ^5^. In the TMEV model, mice exhibit seizures 3-8 days postinfection (DPI), clear the virus by DPI 14, and can develop chronic TLE^6^. These mice also exhibit long-term decreases in seizure threshold that may be indicative of increased neuronal excitability^7^ and exhibit anxiety-like behavior and cognitive impairments^8^. TMEV infection also causes pyramidal cell death in hippocampal CA1^6,9^. It is possible that the neuronal damage and innate immune system response to viral infection in CA1^8,10^ provokes acute seizures^10,11,12,13^, while long-term changes in hippocampal circuitry underlie the chronic seizures^14^. However, the precise regions underlying seizures during the acute phase are unknown.

Transient Recombination in Active Populations (TRAP) labels active cells with a fluorescent marker through a 4-hydroxytamoxifen (4-OHT) dependent, Cre-inducible paradigm^15^. TRAP tagging has been used to map seizure propagation in pentylenetetrazol-induced seizures^16^, cobalt-induced frontal lobe seizures^17^, hypoxic-ischemic-induced seizures^18^, and status epilepticus in mice^19^. However, seizure propagation networks have not been examined in the TMEV model. Given the extensive damage observed in CA1 and CA2 of infected mice, it is unknown how seizures propagate in this model during the acute infection period. Here we investigated seizure-active circuitry on day post-injection 5 (DPI 5) of TMEV infection by quantifying c-Fos driven tdTomato expression in seizing c-Fos-CreERT2 x CAG-tdTomato (TRAP mice; n = 14) and PBS-injected control mice (n =7). The c-Fos promoter ensures only seizure-active cells become ‘trapped’ (marked with tdTomato) while the 4-OHT administration provides temporal specificity. Because the timing of 4-OHT administration influences the quantity of trapped cells, 4-OHT was given at a short (1.5 hr) or later (3 hr) timepoint based on prior investigations^19,20^.

We identified seven saliant regions of interest that expressed high levels of tdTomato both ipsilateral and contralateral to the site of injection following seizures in TMEV-injected TRAP mice. This is consistent with seizures becoming secondarily generalized in this model. The hippocampus, thalamus, lateral septal nuclei, basal ganglia, triangular septal nucleus, fornix, and corpus callosum expressed higher tdTomato levels compared to PBS-injected controls (p < 0.05) in the groups receiving 4-OHT 1.5 hr and 3 hrs post-seizure. However, tdTomato expression was reduced in TMEV-injected mice administered 4-OHT at the 3 hr timepoint compared to the 1.5 hr timepoint (p < 0.05). Damage within CA1 was observed in both groups of TMEV-injected mice. Sex differences in tdTomato expression were not observed between seizing male (n = 7) and female (n = 7). TMEV-injected TRAP mice. Examination of the hippocampus indicated that the subregions with the greatest tdTomato expression were the dentate gyrus (DG), hilus, and CA3. This indicates that seizure generalization must arise from hippocampal outputs independent of CA1 and suggests the DG, hilus, and CA3 as salient regions to target during TMEV infection.

## MATERIALS AND METHODS

### Animals

All experimental procedures were conducted in compliance with the Institutional Animal Care and Use Committee at the University of Utah College of Pharmacy. Mice were housed in a 12:12 light:dark cycle (6:00 AM to 6:00 PM). Soy-free mouse chow (2020X; Inotiv, Madison, WI) and water were available ad libitum. Animals received daily health checks and pain and distress were minimized throughout the experiments. Any mice which lost > 20% of their starting weight for at least 24 hr were removed from the experiment.

Male and female TRAP mice (c-Fos-CreERT2 x CAG-tdTomato) and their WT littermates (11-16 wks) were utilized for these experiments^16,18^. To generate c-Fos-CreERT2 x CAG-tdTomato mice, hemizygous mice expressing Cre-ERT2 under the c-Fos promoter (B6.129(Cg)- Fos^tm1.1(cre/ERT2)^Luo/J; #021882; Jackson Laboratories, Bar Harbor, ME) were crossed with homozygous mice expressing floxxed tdTomato ubiquitously under the Rosa26 locus (B6.Cg-Gt(ROSA)26Sor^tm9(CAG-tdTomato)^Hze/J; #007909; Jackson Laboratories, Bar Harbor, ME). Offspring were all hemizygous for CAG-tdTomato. Offspring positive for Cre-ER were the experimental mice. The original stock was obtained from Jackson Labs (Bar Harbor, ME), and were maintained at the animal husbandry facilities at University of Utah. Mice were selected for treatment conditions in a random order. Treatments were administered by one experimenter. The experimenter was blinded to genotypes until data analysis was completed.

### Experimental procedures

#### TMEV Injection

Adult male and female TRAP mice (11-16 wks) were unilaterally inoculated with 20 uL of the Daniel’s strain^21^ of TMEV (1.67 x 10^7^ p/mL) or PBS via intracranial injection near the midline of the right parietal cortex (28 ga insulin syringe)^22,23^. TMEV-injected mice have spontaneous seizures and experience seizures following gentle handling. Focal seizures may be observed on DPI 2 and 3. However, seizures escalate to generalized-tonic clonic seizures between DPI 4-7^5,22^. During these experiments, mice were handled from DPI 3 – DPI 5 to check for seizures. If at least 60% of a cohort experienced seizures, the animals were tested. On the morning of DPI 5, all mice were handled to induce seizures or recapitulate the conditions leading to a seizure in PBS mice. Seizures were evaluated according to a modified racine scale (0, no seizure; 1, freezing, facial automatisms; 2, head bobbing, urinating, drooling; 3, unilateral forelimb myoclonus; 4, bilateral myoclonus, rearing and falling; 5, wild running, jumping, loss of posture)^23,24,25^.

The mice that seized (and the PBS-injected mice) received an injection of z-4-hydroxytamoxifen (4-OHT, 50 mg/kg) 1.5 hr or 3 hr after handling. 4-OHT binds to ERT2, allows cytoplasmic Cre recombinase to enter the nucleus, which then promotes expression of tdTomato in previously active cells by cleaving lox-p sites^15^. Mice which had seizures during infection but did not seize on DPI 5 were excluded from study. Similarly, mice which displayed no seizures during infection were also excluded, as it is inconclusive if those mice ever seized. Only mice with racine grade 5 seizures were utilized for the studies.

#### 4-Hydroxytamoxifen Preparat*ion and Injection*

4-OHT (Sigma, H6278) is a modulator at estrogen receptors including the mutant estrogen receptor ERT2. It is the metabolically active form of the estrogen receptor antagonist tamoxifen and has a 100X infinity for the estrogen receptor. 4-OHT is cleared faster than tamoxifen by avoiding first-pass metabolism^26^. This modified receptor is used in Cre-inducible paradigms to differentiate cell types and networks based on fluorescent reporters. 4-OHT solutions were made fresh on the day of experimentation by dissolving 4-OHT in 100% EtOH. Aliquots were added to peanut oil to reach a final concentration of 10% EtOH. The 4-OHT was stirred on a 60 C heated plate until dissolved. Injections of 4-hydroxytamoxifen (50 mg/kg) were delivered either 1.5 or 3 hr *i.p.* following a handling-induced seizure. 4-OHT is mostly eliminated in 6-8 hr^27^ and is less effective at trapping by this timepoint^19,27^, therefore the mice were video monitored for 6 hr following 4-OHT to ensure animal health as well as monitor for seizures.

### Tissue processing

#### Intracardiac perfusion

Mice were transcardially perfused with chilled 1x PBS (4C) followed by paraformaldehyde (4%) 7 days after the 4-OHT injections to optimize tdTomato expression. Extracted brains remained in chilled 4% PFA for 1 day and then were cryoprotected in 30% sucrose for 3 days prior to horizontal sectioning on a microtome (20 uM; SM2010R; Leica Microsystems; Wetzlar, Germany). Sections were then mounted for immunohistochemistry.

#### Immunohistochemistry

Briefly, tissue was treated at room temperature with three 1x PBS washes before overnight incubation in a primary antibody solution containing rabbit anti DSred (1:1000; 632496; Takara, San Jose, CA). The tissue was then washed 3x with 1x PBS before being incubated in secondary antibody solution containing Cy3 donkey anti-rabbit (1:500; 712165153; Jackson Immunoresearch, West Grove, PA). The types of seizure active, trapped cells were assessed in sections following an identical protocol. Mouse anti-NeuN (1:1000; MAB377; EMD Millipore, St. Louis, MO) and chicken anti-GFAP (1:1000; ab4674; Abcam, Eugene, OR) were added to the primary antibody solution. Cy5 donkey anti-mouse (1:500; 71517515) and A488 Donkey anti-chicken (1:500; 703545155) from Jackson Immunoresearch were added to the secondary antibody solution. The primary anti-Dsred antibody was used to enhance tdTomato visibility, the NeuN antibody indicated neuron cell bodies, and the GFAP antibody identified astrocytes. All antibodies were diluted in CytoQ ImmunoDiluent & Block Solution (NB307-C; Innovex; Richmond, CA) and 0.05% Triton x-100 to permeabilize the membrane and prevent nonspecific binding. Three brain sections per animal were used for quantification. Regions of interest were split along the longitudinal fissure to indicate structures ipsilateral vs contralateral to the site of injection. Slides were stained in batches containing control and experimental animals. Digital fluorescent images were obtained on a Zeiss Axioscan 7 (Zeiss, Oberkochen, Germany) slide scanner at 10x. Robust trapped tdTomato expression was observed within the hippocampus, so higher magnification (40x) images were captured of the HPC and the CA1, CA3, and the dentate gyrus/hilus. Anatomic landmarks were identified with the aid of a mouse brain atlas^28^. Imaging parameters remained consistent throughout scanning. Resolution was set at 0.32 um pixel size with a 10% stitching overlap.

#### c-Fos Quantification

QuPath is a high-throughput biomarker evaluation tool for digital slides^29^. Three horizontal sections (∼-5.6 mm from bregma) were analyzed from each mouse. The following were chosen as brain regions of interest and annotated: hippocampi, thalamus, lateral septal nuclei, basal ganglia, triangular septal nucleus, fornix, and corpus callosum. The longitudinal fissure delineated whether a structure was ipsilateral or contralateral to the side of injection. A Gaussian smoothing filter was applied to reduce noise and improve quantification. Cells exhibiting Cy3/tdTomato were considered trapped cells and were identified in QuPath using a thresholder based on optical densities. The same classifier was utilized for all sections within each IHC batch. The area of each region and the area of positively labeled cells were then used to calculate the percent area of fluorescence. This was averaged across three sections for each animal to produce the final quantification.

#### Statistical analyses

Threshold for statistical significance was set at p < 0.05 for all comparisons. Normality of the data was assessed via a Shapiro-Wilk normality test. Unpaired two-tail tests (parametric) or Mann Whitney U-tests (nonparametric) were utilized for between-subjects PBS vs TMEV comparisons and paired t-tests (parametric) or Wilcoxon signed rank tests (nonparametric) were utilized for within-subjects ipsilateral vs contralateral comparisons. P-values were adjusted for false discovery rate using the two-stage step-up method by Benjamini, Krieger, and Yekutieli. Post-hoc power analyses were conducted to confirm ≥0.80 β power for experiments (G*Power; Heinrich Heine University Düsseldorf)^30^. Graphpad Prism 7 software (Boston, MA) was utilized for visualizing results and statistics.

#### Data Availability

Raw data was collected at University of Utah. All data, protocols, and scripts are available upon reasonable request to the corresponding author.

## Results

### TMEV-injected TRAP mice injected with 4-OHT 1.5 hr following seizures exhibit elevated tdTomato expression in ipsilateral structures

TMEV-injected mice experience seizures between DPI 3-7 and have intense bilateral generalized tonic clonic seizures DPI 5-7. Seizures robustly increase c-Fos expression through synchronous activation of neural networks. Here, tdTomato was used as a proxy marker for c-Fos expression following a generalized seizure at DPI 5 in our TRAP mice (n = 14) or gentle handling in PBS controls (n = 7). An increase in tdTomato expression was observed in brain sections from TRAP mice when 4-OHT was administered 1.5 hr following a seizure in the ipsilateral hippocampus (Mann-Whitney test), thalamus (unpaired test), lateral septal nucleus (Mann-Whitney test), basal ganglia (Mann-Whitney test), fornix (unpaired t-test), corpus callosum (unpaired t-test), and triangular septal nucleus (Mann-Whitney test). All p values < 0.05 (Fig. 1A,B). Sex differences in tdTomato expression were not observed between seizing male (n = 7) and female (n = 7) TMEV-injected TRAP mice in the ipsilateral hippocampus (unpaired t-test), thalamus (unpaired t-test), lateral septal nuclei (Wilcoxon test), basal ganglia (unpaired t-test), fornix (unpaired t-test), corpus callosum (Wilcoxon test), and triangular septal nucleus (unpaired t-test). All p values > 0.05 (Fig. 1C,D). Means and SEM are presented in Supplemental Table 1 for clarity. Horizontal brain sections immunostained for tdTomato (cyan) (Fig. 1E) indicated that at DPI 5 seizure-active, c-Fos expressing cells are present throughout the regions of interest in TMEV-injected mice (right image) but are absent in PBS-injected mice (left image). The identity of these seizure-active, c-Fos expressing cells is unknown. The hippocampus was a focus due to CA1 damage by TMEV. The ipsilateral hippocampus was magnified (40x) and immunostained for NeuN (neuronal nuclei, magenta) and GFAP (glial fibrillary acidic protein, green), to assess colocalization with tdTomato (cyan) in TMEV-injected mice (Fig. 1F). TdTomato (cyan) was robustly expressed in neuronal cell bodies, axons, and dendrites within the hippocampus (Fig. 1F), and the CA3 (Fig. 1F,i.), CA1 (Fig. 1F,ii.), and the dentate gyrus / hilus (Fig. 1F,iii.) hippocampal subregions. Astrocytes were colocalized with tdTomato expressing cells in the hilus, although there was little colocalization in other hippocampal subregions.

**Figure 1.**
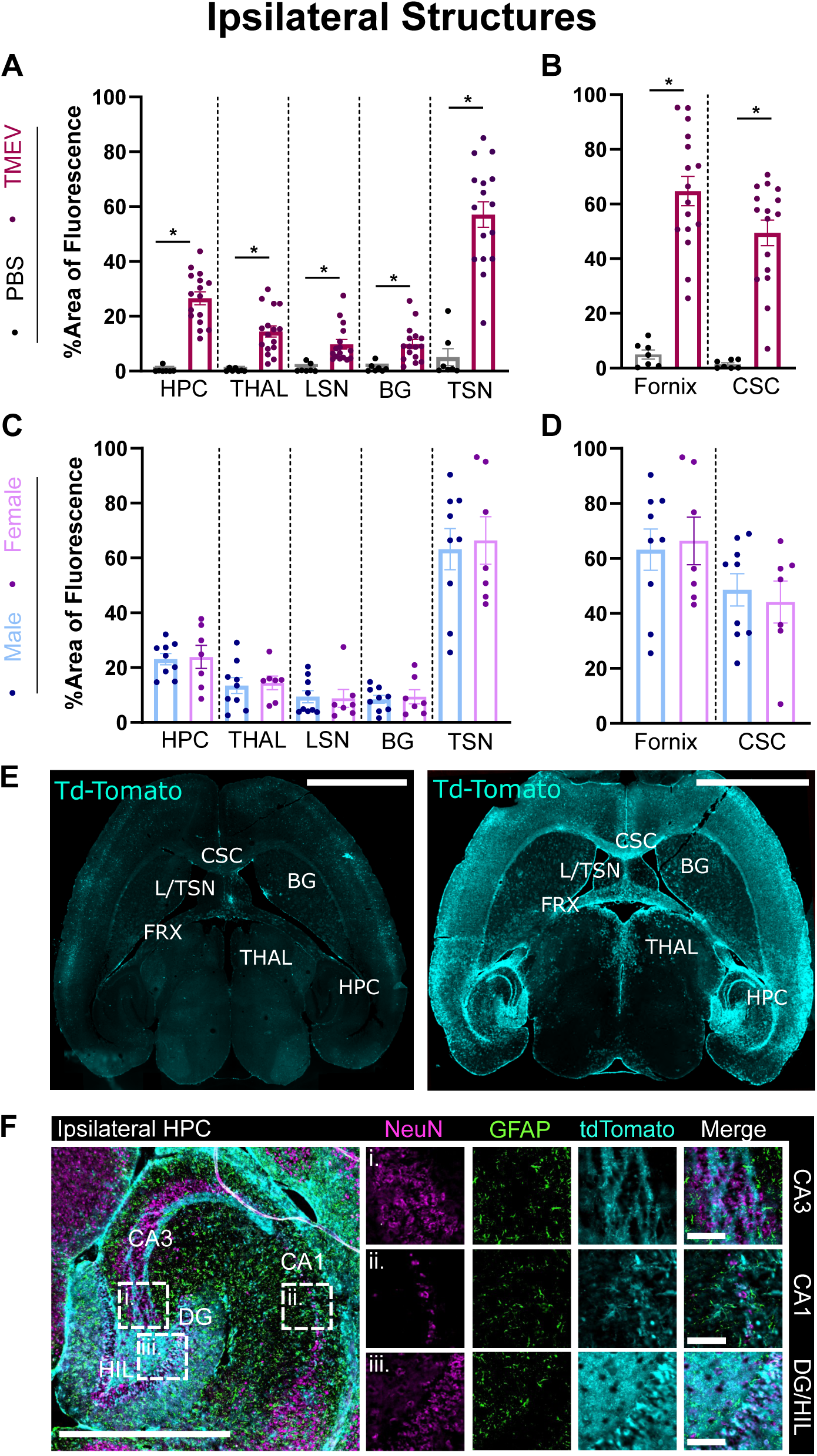
Increased c-Fos activity is observed in ipsilateral structures of TMEV-injected TRAP mice administered 4-OHT 1.5 hr following a seizure on DPI 5. (A,B) Percent area of fluorescence in TMEV-injected TRAP mice (n = 14) vs the non-seizing PBS-injected controls (n = 7) in sites ipsilateral to injection site: hippocampus (*p = 0.0001), thalamus (*p = 0.0005), lateral septal nuclei (*p = 0.0001), basal ganglia (*p = 0.0001), fornix (*p = 0.0001), corpus callosum (*p = 0.0001), and triangular septal nucleus (*p = 0.0001). (C,D) Sex differences in percent area of fluorescence between male (n = 7) and female (n = 7) TRAP animals in structures ipsilateral to injection sites: hippocampus (p = 0.729), thalamus (p = 0.568), lateral septal nuclei (p = 0.383), basal ganglia (p = 0.938), fornix (p = 0.833), corpus callosum (p = 0.902), and triangular septal nucleus (p = 0.454), Bars: black = PBS, purple = TMEV; blue = male, pink = female. Error bars represent Mean ± SEM. Unpaired t-tests or Mann Whitney tests were utilized to analyze data depending on normality. (E) 10x Immunostained horizontal mouse sections demonstrating tdTomato as a proxy for c-Fos expression (cyan) in a PBS-injected (left) and TMEV-injected (right) mouse that received 4-OHT 1.5 hr after a seizure at DPI 5. Scalebar = 2 mm. (F) 40x immunostained horizontal sections of the HPC on the ipsilateral side of TMEV injection (E). Neuronal nuclei and astrocytes were identified via NeuN (magenta) and glial fibrillary acidic protein (GFAP, green) staining and compared to the expression of seizure-active, c-Fos expressing cells (tdTomato, cyan). Boxes indicate insets of CA3 (i.), CA1 (ii.), and DG/HIL (iii.). Pyramidal cell loss due to TMEV infection can be seen in CA1 (ii.). Robust expression of trapped tdTomato (cyan) is evident in CA3 (i.) and the DG/HIL (iii.). Scalebars = 300 um,100 um; BG = basal ganglia; CSC = corpus callosum; GFAP = glial fibrillary acidic protein; HPC = hippocampus; LSN = lateral septal nucleus; NeuN = neuronal nuclei; THAL = thalamus; TSN = triangular septal nucleus.

### TMEV-injected TRAP mice injected with 4-OHT 1.5 hr following seizures also exhibit elevated tdTomato expression in contralateral structures

Generalized seizures propagate to both hemispheres of the brain. Likewise, TMEV spreads from the point of injection (the ipsilateral side of the brain) to the contralateral side of the brain during infection. Thus, we split regions of interest along the longitudinal fissure to determine any differences in c-Fos expression between brain hemispheres. Increased tdTomato expression was also observed in horizontal brain sections obtained from TMEV-injected TRAP mice (n = 14) compared to the non-seizing PBS-injected controls (n = 7) following seizures induced at DPI 5 in the contralateral hippocampus (Mann Whitney test), thalamus (unpaired t-test), lateral septal nucleus (Mann Whitney test), basal ganglia (Mann Whitney test), fornix (unpaired t-test), and corpus callosum, (Mann Whitney test). All p values < 0.05 (Fig. 2A,B). No differences in tdTomato expression were observed when these mice were sub-grouped by sex (7 male and 7 female): contralateral hippocampus (unpaired t-test), thalamus (unpaired t-test), lateral septal nuclei (unpaired t-test), basal ganglia (unpaired t-test), fornix (unpaired t-test), and corpus callosum (unpaired t-test). All p values > 0.05 (Fig. 2C,D). Means and SEM are presented in Supplemental Table 1. The contralateral hippocampus was marked as a saliant site (Fig. 2E) using the immunostaining presented in Fig. 1E. The hippocampus was magnified (40x) in horizontal brain sections immunostained for NeuN (magenta), GFAP (green), and tdTomato (cyan) in TMEV- injected mice after a seizure on DPI 5 (Fig. 2F). As in Fig. 1F, there was robust expression of tdTomato (cyan) in neuronal cell bodies, axons, and dendrites within the HPC (Fig. 2F), and the CA3 (Fig. 2Fi.), CA1 (Fig. 2Fii.), and the dentate gyrus / hilus (Fig. 2Fiii.) hippocampal subregions. No change in colocalization with NeuN or GFAP was noted.

**Figure 2.**
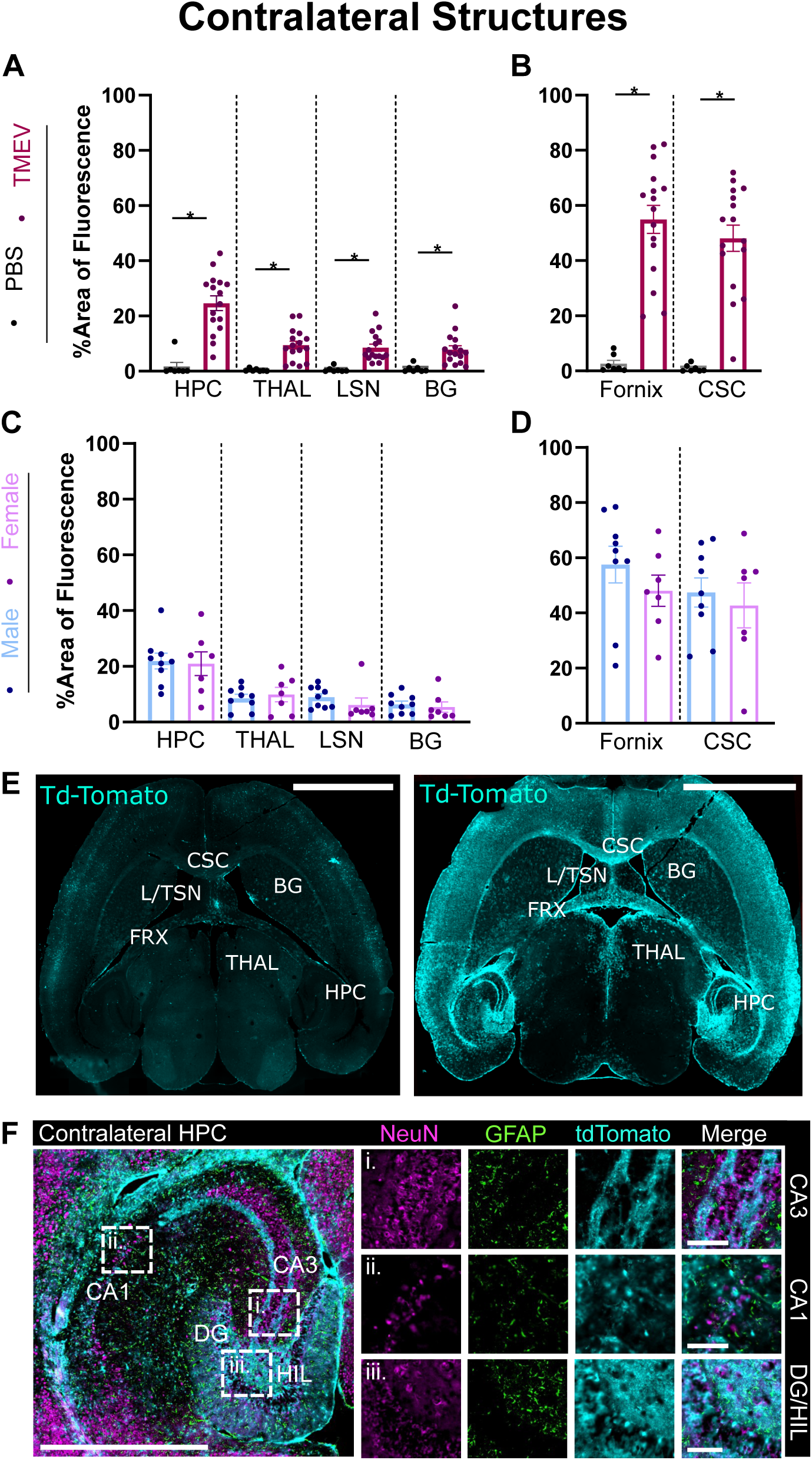
Increased c-Fos activity is observed in contralateral structures of TMEV-injected TRAP mice administered 4-OHT 1.5 hr following a seizure on DPI 5. (A,B) Percent area of fluorescence in TMEV-injected seizing TRAP mice (n = 14) vs the non- seizing PBS-injected controls (n = 7) in sites contralateral to the injection site: hippocampus (*p = 0.0002), thalamus (*p = 0.0005), lateral septal nuclei (*p = 0.0001), basal ganglia (p = 0.0003), fornix (*p = 0.0001), and corpus callosum (*p = 0.0001). (C,D) Sex differences in percent area of fluorescence between male (n = 7) and female (n = 7) animals in structures contralateral to injection sites: hippocampus (p = 0.997), thalamus (p = 0.141), lateral septal nuclei (p = 0.950), basal ganglia (p = 0.304), fornix (p = 0.270), and corpus callosum (p = 0.678). Bars: black = PBS, purple = TMEV; blue = male, pink = female. Error bars represent Mean ± SEM. Unpaired t-tests or Mann Whitney tests were utilized to analyze data depending on normality. (E) 10x Immunostained horizontal mouse sections stained for tdTomato (cyan) as a proxy for c-Fos expression in a PBS-injected (left) and a TMEV-injected TRAP mouse (right) that received 4-OHT 1.5 hr after a seizure at DPI 5. Scalebar = 2 mm. (F) 40x immunostained horizontal mouse sections of the HPC on the contralateral side of TMEV injection (E). Co-expression of seizure- active, c-Fos expressing cells (tdTomato, cyan), NeuN (magenta), and GFAP (green) were assessed in CA3 (i.), CA1 (ii.), and the DG/HIL (iii.). Boxes indicate inset location. Scale bars = 300 um, 100 um; BG = basal ganglia; CSC = corpus callosum; GFAP = glial fibrillary acidic protein; HPC = hippocampus; LSN = lateral septal nucleus; NeuN = neuronal nuclei; THAL = thalamus; TSN = triangular septal nucleus.

### TMEV-injected TRAP mice that receive 4-OHT 1.5 hr following a seizure exhibit no differences in tdTomato expression across ipsilateral vs contralateral structures

Although TMEV was injected in the right (ipsilateral) hemisphere, tdTomato was trapped in both hemispheres. There were no differences in tdTomato between the ipsilateral vs contralateral hippocampus (paired t-test), thalamus (paired t-test), lateral septal nuclei (Wilcoxon test), basal ganglia (paired t-test), fornix (paired t-test) and corpus callosum (paired t-test). All p > 0.05 (n = 14; Fig. 3A-D). This activity across the ipsilateral and contralateral hemispheres is consistent with the propagation of generalized seizures. Similarly, there were only minor changes in tdTomato expression between hemispheres of PBS-injected mice (n = 7; Fig. 3C,D): hippocampus (Wilcoxon test), thalamus (paired t-test), basal ganglia (Wilcoxon test), and corpus callosum (Wilcoxon test). All p > 0.05. There were small decreases in tdTomato expression across the lateral septal nuclei (Wilcoxon test) and fornix (Wilcoxon test; p < 0.05).

**Figure 3.**
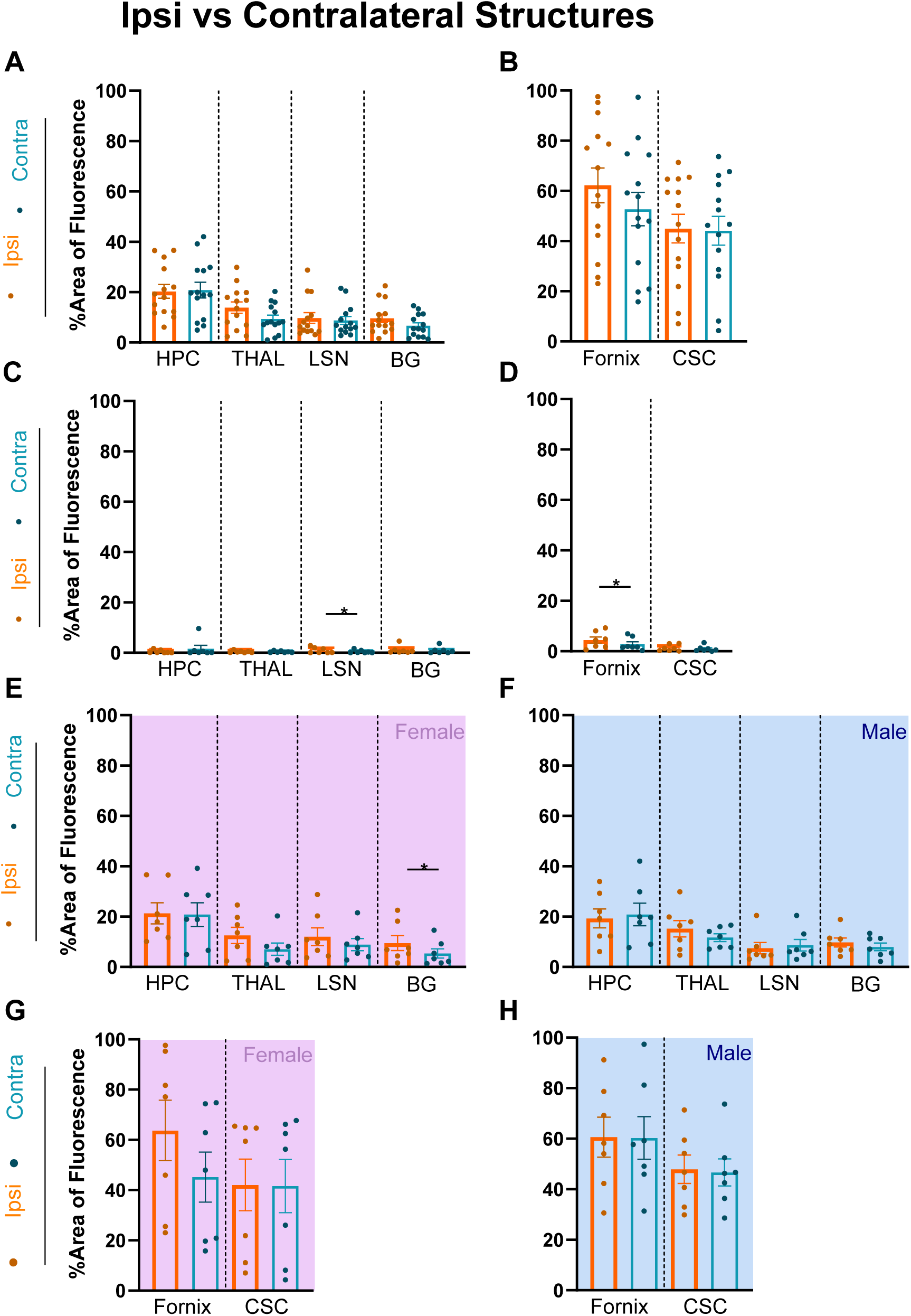
c-Fos expression is similar between ipsilateral and contralateral structures in TMEV-injected TRAP mice injected with 4-OHT 1.5 hr post seizure on DPI 5. (A,B) tdTomato area of fluorescence between structures ipsilateral and contralateral to the TMEV injection site in seizing TRAP mice (n = 14): hippocampus (p = 0.751), thalamus (p = 0.080), lateral septal nuclei (p = 0.542), basal ganglia (p = 0.062), fornix (p = 0.129), and corpus callosum (p = 0.579). (C,D) tdTomato expression between structures ipsilateral and contralateral to the TMEV injection site in PBS-injected non-seizing controls. Hippocampus (p = 0.469), thalamus (p = 0.710), lateral septal nuclei (*p = 0.031), basal ganglia (p = 0.219), fornix (*p = 0.016), and corpus callosum (p = 0.813). (E-H) Percent area of fluorescence between ipsilateral and contralateral structures within sexes of seizing TMEV-injected TRAP mice. Male (n = 7): hippocampus (p = 0.517), thalamus (p = 0.350), lateral septal nuclei (p = 0.578), basal ganglia (p = 0.516), fornix (p = 0.935), and corpus callosum (p = 0.681). Female (n = 7): hippocampus (p = 0.854), thalamus (p = 0.163), lateral septal nuclei (p = 0.107), basal ganglia (*p = 0.023), fornix (p = 0.115), and corpus callosum (p = 0.813). Bars: orange = ipsilateral, blue = contralateral. Error bars represent Mean ± SEM. Ipsi = ipsilateral, contra = contralateral. Paired t-tests or Wilcoxon tests were used to analyze the within subject comparisons depending on data normality.

No sex differences in tdTomato expression were observed between ipsilateral and contralateral regions of interest within male (n = 7) and female (n = 7) TMEV-injected TRAP mice (Fig. 3E-H). Area of tdTomato fluorescence was comparable (p > 0.05) in the hippocampus (paired t-test), thalamus (paired t-test), lateral septal nuclei (Wilcoxon test), basal ganglia (paired t-test), fornix (paired t-test), and corpus callosum (paired t-test; p < 0.05) of male TRAP mice (Fig. 3F,H). Comparisons between ipsilateral and contralateral regions in female TMEV-injected TRAP mice revealed no changes (p > 0.05) within the hippocampus (paired t-test), thalamus (paired t-test), lateral septal nuclei (paired t-test), basal ganglia (paired t-test), fornix (paired t-test), and corpus callosum (Wilcoxon test; Fig. 3E,G). All means and SEM are presented in Supplemental Table 1.

### TMEV-injected TRAP mice administered 4-OHT 3 hr after seizures exhibit differences in tdTomato expression in ipsilateral and contralateral structures

The temporal specificity of cFos-TRAP depends on the timing of 4-OHT injection. We utilized two 4-OHT injection timepoints (1.5 and 3 hr post-seizure) to ensure seizure-active cells were optimally captured^19,31^. Like the group administered 4-OHT 1.5 hr post-seizure, increased tdTomato expression was observed in the ipsilateral structures of TMEV-injected (n = 13) and PBS-injected (n = 7) TRAP mice. In the group that received 4-OHT 3 hr post-seizure, tdTomato was elevated in the ipsilateral hippocampus (Mann-Whitney test), thalamus (unpaired t-test), lateral septal nucleus (Mann-Whitney test), basal ganglia (Mann-Whitney test), fornix (unpaired t- test), corpus callosum (unpaired t-test), and triangular septal nucleus (unpaired t-test; Fig. 4A,B) when compared to PBS controls. The expression of tdTomato was likewise elevated in the contralateral hippocampus (Mann Whitney test), thalamus (Mann Whitney test), lateral septal nucleus (Mann Whitney test), basal ganglia (Mann Whitney test), fornix (Mann Whitney test), and corpus callosum (unpaired t-test; Fig. 4C,D). All p < 0.05.

**Figure 4.**
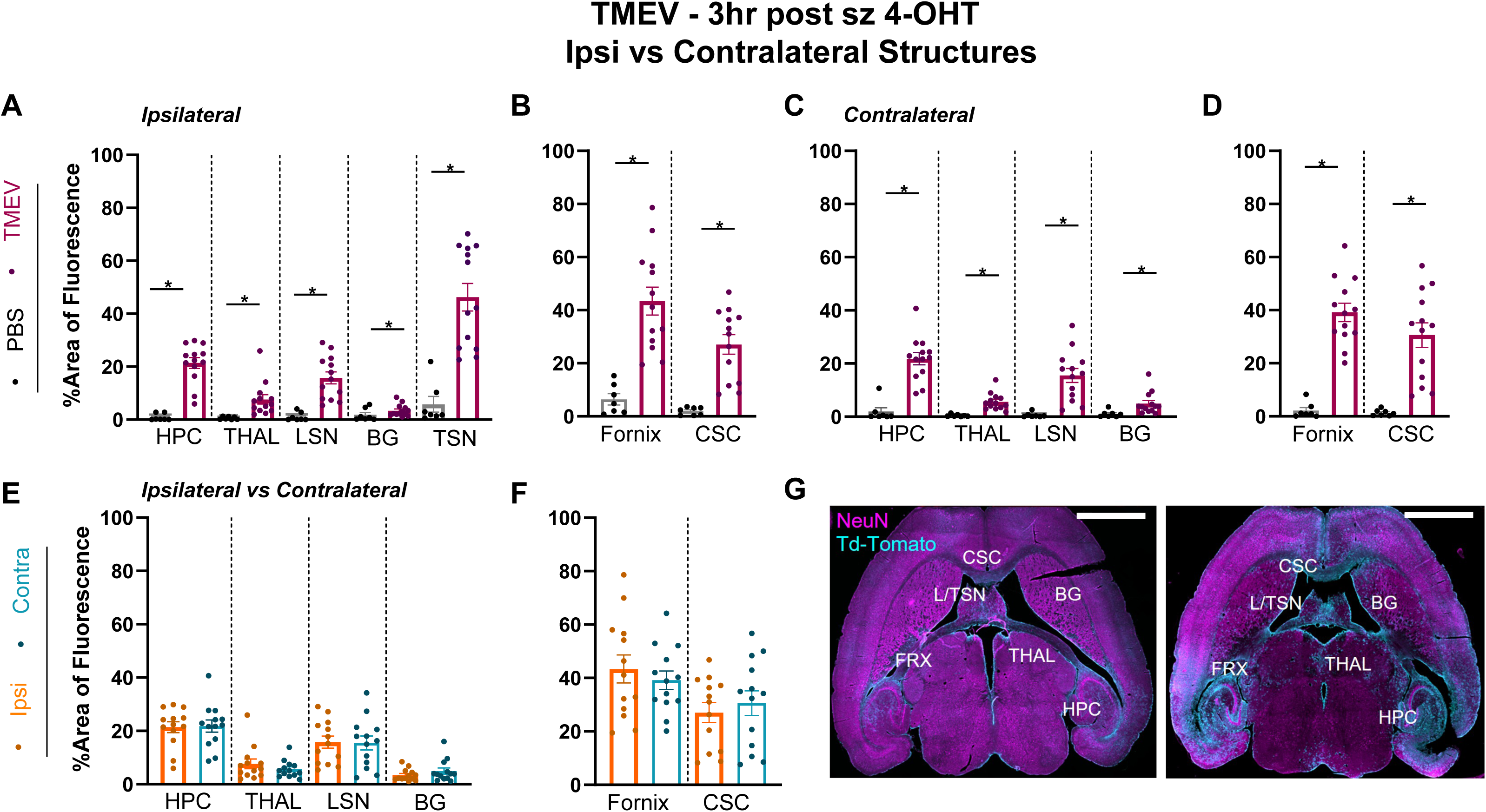
Increased c-Fos activity is observed in ipsilateral structures of TMEV-injected TRAP mice administered 4-OHT 3 hr following a seizure on DPI 5 (A,B) Percent area of fluorescence of tdTomato in TMEV-injected seizing TRAP mice (n = 13) and the PBS-injected controls (n = 7) in sites ipsilateral to injection site: hippocampus (*p = 0.0001), thalamus (*p = 0.001), lateral septal nuclei (*p = 0.0001), basal ganglia (*p = 0.002), fornix (*p = 0.0001), corpus callosum (*p = 0.0001), and triangular septal nucleus (*p = 0.0001). Bars: black = PBS, purple = TMEV. Error bars represent Mean ± SEM. (C,D) tdTomato expression in TMEV-injected seizing TRAP mice (n = 13) and PBS-injected controls (n = 7) in sites contralateral to injection site: hippocampus (*p = 0.0001), thalamus (*p = 0.0001), lateral septal nuclei (*p = 0.0001), basal ganglia (*p = 0.0005), fornix (*p = 0.0001), corpus callosum (*p = 0.0001). Bars: black = PBS, purple = TMEV; Error bars represent Mean ± SEM. (E,F) TdTomato expression between ipsilateral and contralateral structures in seizing TRAP mice administered 4- OHT 3 hr after their seizure. Hippocampus (p = 0.862), thalamus (p = 0.455), lateral septal nuclei (p = 0.579), basal ganglia (p = 0.191), fornix (p = 0.372), and corpus callosum (p = 0.372). Bars: orange = ipsilateral, blue = contralateral. (G) Immunostained horizontal mouse sections demonstrating tdTomato as a proxy for c-Fos expression (cyan) and NeuN (magenta) in a PBS-injected (left) and TMEV-injected (right) mouse. Scalebar = 2 mm; HPC = hippocampus; THAL = thalamus; BG = basal ganglia; L/TSN = lateral/triangular septal nucleus; CSC = corpus callosum. (A-D) Unpaired t-tests or Mann Whitney tests were utilized to analyze data depending on normality. (E,F) Paired t-tests or Wilcoxon tests were used to analyze the within subject comparisons depending on data normality.

Next tdTomato fluorescence was examined between ipsilateral and contralateral regions within the TRAP mice (n = 13; Fig. 4E,F) that were administered 4-OHT 3 hr following seizures induced at DPI 5. No differences in tdTomato expression were noted between the ipsilateral and contralateral hippocampus (paired t-test), thalamus (Wilcoxon test), lateral septal nuclei (paired t- test), basal ganglia (Wilcoxon test), fornix (paired t-test), and corpus callosum (paired t-test). All p > 0.05. Means and SEM are presented in Supplemental Table 2 for clarity.

### TMEV-injected TRAP mice given 4-OHT 3 hr after a seizure have reduced tdTomato expression in some structures compared to TMEV-injected TRAP mice administered 4-OHT 1.5 hr after a seizure

The timing of 4-OHT injection determines the window when active cells may be trapped. When we examined tdTomato fluorescence between the 1.5 hr vs the 3 hr post-seizure 4-OHT injections, there was a decrease in fluorescence across the ipsilateral thalamus (Mann Whitney test), lateral septal nuclei (unpaired t-test), basal ganglia (Mann Whitney test), fornix (unpaired t- test), and corpus callosum (unpaired t-test; p < 0.05; Fig. 5A,B). Uniquely, the lateral septal nuclei (Mann Whitney test) displayed elevated tdTomato expression in the 3 hr compared to the 1.5 hr 4-OHT timepoint (p > 0.05).

**Figure 5.**
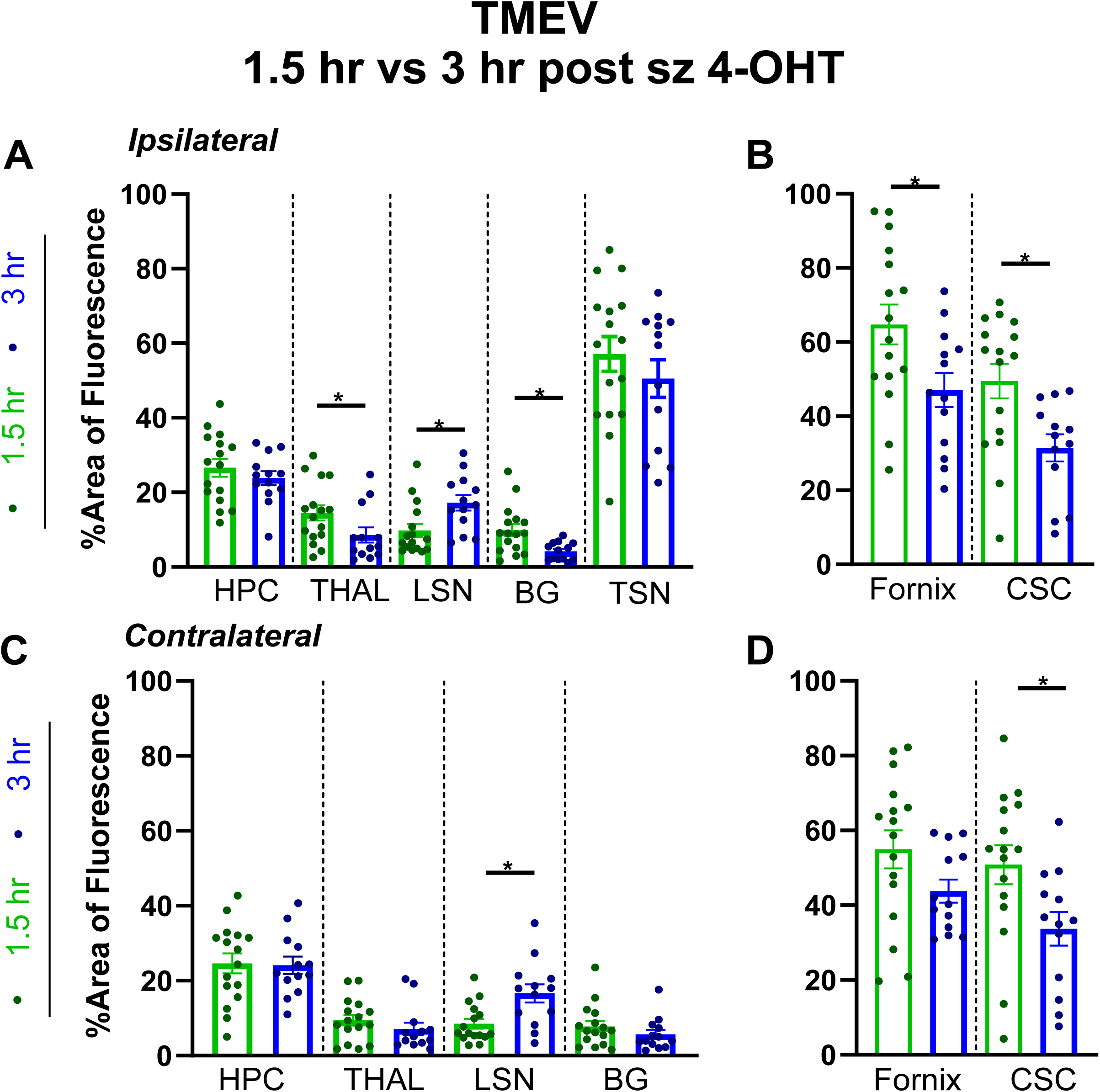
TMEV-injected TRAP mice given 4-OHT 3 hr after a seizure have lower tdTomato expression compared to TMEV-injected TRAP mice administered 4-OHT 1.5 hr after a seizure. (A,B) Percent tdTomato expression in sites ipsilateral to injection site in TMEV-injected TRAP mice that received 4-OHT either 1.5 hr (n = 14) or 3 hr (n =13) after a seizure: hippocampus (p = 0.0.382), thalamus (*p = 0.032), lateral septal nuclei (*p = 0.007), basal ganglia (*p = 0.0007), fornix (*p = 0.023), corpus callosum (*p = 0.007), and triangular septal nucleus (p = 0.351). (C,D) Percent tdTomato expression in sites contralateral to the injection site of TMEV-injected TRAP mice that received 4-OHT either 1.5 hr (n = 14) or 3 hr (n =13) after a seizure: hippocampus (p = 0.892), thalamus (p = 0.215), lateral septal nuclei (*p = 0.004), basal ganglia (p = 0.250), fornix (p = 0.085), corpus callosum (*p = 0.022). Bars: green = 1.5 hr, blue = 3 hr. Error bars represent Mean ± SEM. (A-D) Unpaired t-tests or Mann Whitney tests were utilized to analyze data depending on normality.

In the mice administered 4-OHT 3 hrs post-seizure, the ipsilateral structures had decreased tdTomato expression compared to mice receiving the 1.5 hr 4-OHT injection. However, this effect was blunted contralaterally. Neither the hippocampus (Mann Whitney test), thalamus (Mann Whitney test), lateral septal nuclei (Mann Whitney test), basal ganglia (Mann Whitney test), fornix (Mann Whitney test), and the corpus callosum Mann Whitney test) exhibited differences in tdTomato fluorescence. All p > 0.05. There was increased tdTomato expression in the contralateral LSN (unpaired t-test; p < 0.05) as it was in the ipsilateral LSN. All means and SEM are presented in Supplemental Table 2 for clarity.

## Discussion

Over 65 million patients globally are diagnosed with epilepsy^32^. Viral encephalitis is a common cause of acquired epilepsy^2^. Prior to the present study, there was little information on the neural circuits underlying seizures during the acute and post-viral chronic phase of TMEV infection. TMEV directly infects neurons, causing cell death and hippocampal network disruption, which has been implicated in TMEV-induced seizures^14,33,34,35^. At the height of acute seizure activity (DPI ∼5) and into the chronic phase, there is loss of CA1 pyramidal cell neurons, increased microglia reactivity in the hippocampus and cortex, and release of pro-inflammatory cytokines ^6,11^. The combination of neuroinflammation, direct neuronal damage by TMEV, and the innate immune response creates a pro-epileptogenic environment in the infected mouse brain^7^. Seizures themselves can cause inflammation, therefore a positive-feedback loop may exist between active infection and continued seizures.

Here we identified neuronal networks involved in acute, infection-induced seizures in TMEV- injected mice using TRAP. Mice received 4-OHT either 1.5 hr or 3 hrs after a handling-induced seizure on DPI 5. We examined several regions of interest including the hippocampus, thalamus, basal ganglia, lateral septal nuclei, triangular septal nucleus, fornix, and corpus callosum. These regions expressed high levels of tdTomato following seizures at the two points when 4-OHT was administered (Fig. 1A,B; Fig. 2A,B; Fig. 4A-D; Fig. 5A-D). There were no hemispheric differences (Fig. 3A,B; Fig. 4 E,F) in tdTomato expression. No sex differences in tdTomato expression were observed between male and female TMEV-injected TRAP mice (Fig. 1C,D; Fig. 2C,D).

The hippocampus is a salient region of interest because it is implicated in over 80% of TLE cases ^36^. Projections from CA1 to the subiculum and CA3 to fornix are major outputs from the hippocampus^37,38^. CA1 is damaged in our TMEV-injected mice and could prevent hippocampal output. However, increased tdTomato expression was observed within the ipsilateral and contralateral DG, hilus, and fornix of the TMEV-injected mice (Fig. 1A,B,F; Fig. 2A,B,F). This suggests seizures are generalizing to the contralateral hippocampus via another circuit. Glutamatergic projections to CA3 from DG granule cells cross to the contralateral hippocampus along the associational commissure^39,40^. Mossy cells from the hilus and DG likewise project to the contralateral hilus and DG through the associational commissure^37^. Recurrent excitatory projections within CA3^41^ may amplify changes in synaptic activity and predispose the TMEV- injected mice to further seizures. Alternatively, the seizures may be crossing contralaterally through the alveus. The alveus is formed by the axons of pyramidal cells that then project to the fimbria, onto the fornix, and sends outputs to the mamillary bodies and anterior thalamus^42^.

We have identified a seizure-active hippocampal circuit in the TMEV model using the c-Fos TRAP paradigm. However, the precise cell types involved in seizure circuits induced by TMEV are unknown. TMEV infection damages pyramidal CA1 neurons and induces activation of astrocytes, microglia, and NG2^6,43,44^. Neurons translate c-Fos protein following fluctuations in positive ions (e,g, K^+^, Ca^2+^). This can include depolarization, injury, or repeated glutamatergic signaling^45^. One possibility is that reactive astrocytes could also be driving c-Fos in our TRAP mice, as cell division and increases in TNF-α, IL-1β, and IFN-γ contribute to glial c-Fos expression^45,46^. In these experiments, c-Fos-driven tdTomato was prevalent throughout the hippocampus, especially CA3, the dentate gyrus, and hilus(Fig. 1E; Fig. 2E). Mossy fiber axons could be observed projecting toward CA3 pyramidal cells (Fig. 1F; Fig. 2F). Colocalization between the c-Fos expressing cells, neurons, and astrocytes (Fig. 1F; 2F) indicated low colocalization of tdTomato with astrocytes. However, overlap of tdTomato, astrocytes, and neurons is present within the hilus. The hilus contains excitatory mossy cells and several types of inhibitory interneurons. It is possible that inhibitory interneuron axons comprise the robust tdTomato staining in the hilus.

Future experiments will utilize spatial transcriptomics to more accurately assay the cell populations within the DG, hilus, and CA3 that are active during seizures in TMEV-injected mice. Through these experiments and future studies, we hope to find avenues to manipulate discrete, seizure active regions and circuits in TMEV-injected animals and reduce seizure incidence. This could ultimately inform treatments for individuals with severe viral infections

## Abbreviations

4-OHT, z-4-hydroxytamoxifen; BG, basal ganglia; Contra, contralateral; CSC, corpus callosum; DG, dentate gyrus; DPI, days post-injection; ERT2, mutant estrogen receptor 2; FRX, fornix; GFAP, glial fibrillary acidic protein; HIL, hilus; HPC, hippocampus; hr, hour; IHC, immunohistochemistry; intracranial, i.c.; Ipsi, ipsilateral; LSN, lateral septal nuclei; NeuN, neuronal nuclei; PBS, phosphate buffered saline; ROI, region of interest; THAL, thalamus; TLE, temporal lobe epilepsy; TMEV, Theiler’s murine encephalomyelitis virus; TRAP, transient recombination in active populations; TSN, triangular septal nucleus; s, second; sz, seizure; WT, wildtype

## BIBLIOGRAPHY

1. Singhi P. Infectious causes of seizures and epilepsy in the developing world. Dev Med Child Neurol 53, 600–609 (2011).

2. Zhang P, Yang Y, Zou J, Yang X, Liu Q, Chen Y. Seizures and epilepsy secondary to viral infection in the central nervous system. Acta Epileptologica 2, 12 (2020).

3. Elza Márcia TY, et al. Common infectious and parasitic diseases as a cause of seizures: geographic distribution and contribution to the burden of epilepsy. Epileptic Disorders 24, 994–1019 (2022).

4. Bonello M, Michael BD, Solomon T. Infective Causes of Epilepsy. Semin Neurol 35, 235–244 (2015).

5. Libbey JE, et al. Seizures following picornavirus infection. Epilepsia 49, 1066–1074 (2008).

6. Stewart K-AA, Wilcox KS, Fujinami RS, White HS. Development of Postinfection Epilepsy After Theiler’s Virus Infection of C57BL/6 Mice. Journal of Neuropathology & Experimental Neurology 69, 1210–1219 (2010).

7. Stewart K-AA, Wilcox KS, Fujinami RS, White HS. Theiler’s virus infection chronically alters seizure susceptibility. Epilepsia 51, 1418–1428 (2010).

8. Umpierre AD, et al. Impaired cognitive ability and anxiety-like behavior following acute seizures in the Theiler’s virus model of temporal lobe epilepsy. Neurobiology of Disease 64, 98–106 (2014).

9. Zhan J, et al. Diffusion Basis Spectrum and Diffusion Tensor Imaging Detect Hippocampal Inflammation and Dendritic Injury in a Virus-Induced Mouse Model of Epilepsy. Frontiers in Neuroscience 12, (2018).

10. Kirkman NJ, Libbey JE, Wilcox KS, White HS, Fujinami RS. Innate but not adaptive immune responses contribute to behavioral seizures following viral infection. Epilepsia 51, 454–464 (2010).

11. Vezzani A, et al. Infections, inflammation and epilepsy. Acta Neuropathologica 131, 211–234 (2016).

12. DePaula-Silva AB. The Contribution of Microglia and Brain-Infiltrating Macrophages to the Pathogenesis of Neuroinflammatory and Neurodegenerative Diseases during TMEV Infection of the Central Nervous System. Viruses 16, 119 (2024).

13. DePaula-Silva AB, Bell LA, Wallis GJ, Wilcox KS. Inflammation Unleashed in Viral-Induced Epileptogenesis. Epilepsy Currents 21, 433–440 (2021).

14. Smeal RM, Fujinami R, White HS, Wilcox KS. Decrease in CA3 inhibitory network activity during Theiler’s virus encephalitis. Neuroscience Letters 609, 210–215 (2015).

15. Guenthner Casey J, Miyamichi K, Yang Helen H, Heller HC, Luo L. Permanent Genetic Access to Transiently Active Neurons via TRAP: Targeted Recombination in Active Populations. Neuron 78, 773–784 (2013).

16. Naik AA, Sun H, Williams CL, Weller DS, Julius Zhu J, Kapur J. Mechanism of seizure-induced retrograde amnesia. Progress in Neurobiology 200, 101984 (2021).

17. Singh T, Batabyal T, Kapur J. Neuronal circuits sustaining neocortical-injury-induced status epilepticus. Neurobiology of Disease 165, 105633 (2022).

18. Burnsed J, Skwarzyńska D, Wagley PK, Isbell L, Kapur J. Neuronal Circuit Activity during Neonatal Hypoxic–Ischemic Seizures in Mice. Annals of Neurology 86, 927–938 (2019).

19. Dabrowska N, et al. Parallel pathways of seizure generalization. Brain 142, 2336–2351 (2019).

20. Joshi S, Williamson J, Moosa S, Kapur J. Progesterone Receptor Activation Regulates Sensory Sensitivity and Migraine Susceptibility. The Journal of Pain 25, 642–658 (2024).

21. Daniels JB, Pappenheimer AM, Richardson S. Observations on encephalomyelitis of mice (DA strain). J Exp Med 96, 517–530 (1952).

22. Batot G, Metcalf CS, Bell LA, Pauletti A, Wilcox KS, Bröer S. A Model for Epilepsy of Infectious Etiology using Theiler’s Murine Encephalomyelitis Virus. J Vis Exp, (2022).

23. Metcalf CS, et al. Screening of prototype antiseizure and anti-inflammatory compounds in the Theiler’s murine encephalomyelitis virus model of epilepsy. Epilepsia Open 7, 46–58 (2022).

24. Racine RJ. Modification of seizure activity by electrical stimulation. II. Motor seizure. Electroencephalogr Clin Neurophysiol 32, 281-294 (1972).

25. Goddard GV, McIntyre DC, Leech CK. A permanent change in brain function resulting from daily electrical stimulation. Experimental Neurology 25, 295–330 (1969).

26. Lim YC, Desta Z, Flockhart DA, Skaar TC. Endoxifen (4-hydroxy-N-desmethyl-tamoxifen) has anti-estrogenic effects in breast cancer cells with potency similar to 4-hydroxy- tamoxifen. Cancer Chemother Pharmacol 55, 471–478 (2005).

27. Reid JM, et al. Pharmacokinetics of endoxifen and tamoxifen in female mice: implications for comparative in vivo activity studies. Cancer Chemotherapy and Pharmacology 74, 1271–1278 (2014).

28. Franklin KBJ, Paxinos G. Paxinos and Franklin’s The mouse brain in stereotaxic coordinates (2013).

29. Bankhead P, et al. QuPath: Open source software for digital pathology image analysis. Scientific Reports 7, 16878 (2017).

30. Faul F, Erdfelder E, Lang A-G, Buchner A. G*Power 3: A flexible statistical power analysis program for the social, behavioral, and biomedical sciences. Behavior Research Methods 39, 175–191 (2007).

31. Brodovskaya A, Shiono S, Sun C, Perez-Reyes E, Kapur J. Preferential superficial cortical layer activation during seizure propagation. Epilepsia 66, 929–941 (2025).

32. Milligan TA. Epilepsy: A Clinical Overview. The American Journal of Medicine 134, 840–847 (2021).

33. Patel DC, et al. Hippocampal TNFα Signaling Contributes to Seizure Generation in an Infection-Induced Mouse Model of Limbic Epilepsy. eNeuro 4, (2017).

34. Gerhauser I, Hansmann F, Ciurkiewicz M, Löscher W, Beineke A. Facets of Theiler’s Murine Encephalomyelitis Virus-Induced Diseases: An Update (2019).

35. Gibbons MB, Smeal RM, Takahashi DK, Vargas JR, Wilcox KS. Contributions of astrocytes to epileptogenesis following status epilepticus: opportunities for preventive therapy? Neurochemistry international 63, 660–669 (2013).

36. Tatum WOIV. Mesial Temporal Lobe Epilepsy. Journal of Clinical Neurophysiology 29, (2012).

37. Zappone CA. Hippocampal associational and commissural pathways: Anatomical and electrophysiological studies in the rat.). The University of Arizona. (2004).

38. Cherubini E, Miles R. The CA3 region of the hippocampus: how is it? What is it for? How does it do it? Front Cell Neurosci 9, 19 (2015).

39. Kesner R. Neurobiological foundations of an attribute model of memory. Comparative Cognition & Behavior Reviews 8, 29–59 (2013).

40. Amaral DG, Scharfman HE, Lavenex P. The dentate gyrus: fundamental neuroanatomical organization (dentate gyrus for dummies). Prog Brain Res 163, 3–22 (2007).

41. Gunn BG, Pruess BS, Gall CM, Lynch G. Input/Output Relationships for the Primary Hippocampal Circuit. The Journal of Neuroscience 45, e0130242024 (2025).

42. Chauhan P, Jethwa K, Rathawa A, Chauhan G, Mehra S. The Anatomy of the Hippocampus. In: Cerebral Ischemia (ed Pluta R). Exon Publications

43. Loewen JL, Barker-Haliski ML, Dahle EJ, White HS, Wilcox KS. Neuronal Injury, Gliosis, and Glial Proliferation in Two Models of Temporal Lobe Epilepsy. Journal of Neuropathology & Experimental Neurology 75, 366–378 (2016).

44. Bell LA, Wallis GJ, Wilcox KS. Reactivity and increased proliferation of NG2 cells following central nervous system infection with Theiler’s murine encephalomyelitis virus. Journal of Neuroinflammation 17, 369 (2020).

45. Cruz-Mendoza F, Jauregui-Huerta F, Aguilar-Delgadillo A, García-Estrada J, Luquin S. Immediate Early Gene c-fos in the Brain: Focus on Glial Cells. Brain Sci 12, (2022).

46. Lara Aparicio SY, et al. Current Opinion on the Use of c-Fos in Neuroscience. NeuroSci 3, 687–702 (2022).

