## Supplemental Table 1 for "Seizure Circuit Activity in the Theiler’s Murine Encephalomyelitis Virus Model of Infection-induced Epilepsy Using Transient Recombination in Active Populations"

### 1.5 hr 4-OHT:

#### PBS VS TMEV (% area of fluorescence)

| | | HPC | THAL | LSN | BG | FRN | CSC | TSN |
|---|---|---|---|---|---|---|---|---|
| **IPSI:** | PBS | 0.467 ± 0.253 | 0.431 ± 0.159 | 0.984 ± 0.448 | 1.133 ± 0.599 | 4.433 ± 1.248 | 1.327 ± 0.485 | 5.354 ± 2.971 |
| | TMEV | 20.320 ± 2.717 | 13.890 ± 2.231 | 9.739 ± 2.095 | 9.626 ± 1.656 | 62.190 ± 6.927 | 44.980 ± 5.699 | 55.200 ± 6.098 |
| | P VALUE | *0.0001** | *0.0005** | *0.0001** | *0.0001** | *0.0001** | *0.0001** | *0.0001** |
| | | *Mann Whitney U* | *Unpaired t-test* | *Mann Whitney U* | *Mann Whitney U* | *Unpaired t-test* | *Unpaired t-test* | *Mann Whitney U* |
| | | | *t = 4.208, df = 19* | | | *t = 5.798, df = 19* | *t = 5.342, df = 19* | |
| **CONTRA:** | | | | | | | | |
| | PBS | 1.637 ± 1.339 | 0.387 ± 0.134 | 0.442 ± 0.251 | 0.947 ± 0.486 | 2.875 ± 0.956 | 1.025 ± 0.438 | |
| | TMEV | 20.850 ± 3.122 | 9.370 ± 1.543 | 8.782 ± 1.590 | 6.680 ± 1.218 | 52.730 ± 6.603 | 44.140 ± 5.742 | |
| | P VALUE | *0.0002** | *0.0005** | *0.0001** | *0.0003** | *0.0001** | *0.0001** | |
| | | *Mann Whitney U* | *Unpaired t-test* | *Mann Whitney U* | *Mann Whitney U* | *Mann Whitney U* | *Mann Whitney U* | |
| | | | *t = 4.157, df = 19* | | | | | |

| | | HPC | THAL | LSN | BG | FRN | CSC |
|---|---|---|---|---|---|---|---|
| **TMEV:** | IPSI | 20.320 ± 2.717 | 13.890 ± 2.231 | 9.739 ± 2.095 | 9.626 ± 1.656 | 62.190 ± 6.927 | 44.980 ± 5.699 |
| | CONTRA | 20.850 ± 3.122 | 9.370 ± 1.543 | 8.782 ± 1.590 | 6.680 ± 1.218 | 52.730 ± 6.603 | 44.140 ± 5.742 |
| | P VALUE | *0.751* | *0.080* | *0.542* | *0.062* | *0.129* | *0.579* |
| | | *Paired t-test* | *Paired t-test* | *Wilcoxon* | *Paired t-test* | *Paired t-test* | *Paired t-test* |
| | | *t = 0.324, df = 13* | *t = 1.901, df = 13* | | *t = 2.046, df = 13* | *t = 1.623., df = 13* | *t = 0.579, df = 13* |
| **PBS:** | IPSI | 0.467 ± 0.253 | 0.431 ± 0.159 | 0.984 ± 0.448 | 1.133 ± 0.599 | 4.433 ± 1.248 | 1.327 ± 0.485 |
| | CONTRA | 1.637 ± 1.339 | 0.387 ± 0.134 | 0.442 ± 0.251 | 0.947 ± 0.486 | 2.875 ± 0.956 | 1.025 ± 0.438 |
| | P VALUE | *0.469* | *0.710* | *0.031** | *0.219* | *0.016** | *0.813* |
| | | *Wilcoxon* | *Paired t-test* | *Wilcoxon* | *Wilcoxon* | *Wilcoxon* | *Wilcoxon* |
| | | | *t = 390, df = 6* | | | | |

#### SEX DIFFERENCES (% area of fluorescence)

| | | HPC | THAL | LSN | BG | FRN | CSC | TSN |
|---|---|---|---|---|---|---|---|---|
| **IPSI:** | MALE | 19.320 ± 3.761 | 15.230 ± 3.246 | 7.450 ± 2.250 | 9.763 ± 1.809 | 60.630 ± 7.904 | 47.900 ± 5.611 | 60.000 ± 7.711 |
| | FEMALE | 21.320 ± 4.186 | 12.540 ± 3.229 | 12.030 ± 3.494 | 9.488 ± 2.934 | 63.740 ± 8.030 | 42.070 ± 10.320 | 50.410 ± 9.697 |
| | P VALUE | *0.729* | *0.568* | *0.383* | *0.938* | *0.833* | *0.902* | *0.454* |
| | | *Unpaired t-test* | *Unpaired t-test* | *Wilcoxon* | *Unpaired t-test* | *Unpaired t-test* | *Wilcoxon* | *Unpaired t-test* |
| | | *t = 0.355, df = 12* | *t = 0.588, df = 12* | | *t = 0.080; df = 12* | *t = 0.216; df = 12* | | *t = 0.774, df = 12* |
| **CONTRA:** | | | | | | | | |
| | MALE | 20.860 ± 4.509 | 11.670 ± 1.546 | 8.675 ± 2.289 | 7.979 ± 1.547 | 60.270 ± 8.431 | 46.660 ± 5.379 | |
| | FEMALE | 20.840 ± 4.680 | 7.068 ± 2.482 | 8.889 ± 2.390 | 5.380 ± 1.862 | 45.190 ± 9.944 | 41.610 ± 10.570 | |
| | P VALUE | *0.997* | *0.141* | *0.950* | *0.304* | *0.270* | *0.678* | |
| | | *Unpaired t-test* | *Unpaired t-test* | *Unpaired t-test* | *Unpaired t-test* | *Unpaired t-test* | *Unpaired t-test* | |
| | | *t = 0.004, df = 12* | *t = 1.575, df = 12* | *t = 0.065, df = 12* | *t = 1.07, df = 12* | *t = 1.160, df = 12* | *t = 0.426; df = 12* | |

| | | HPC | THAL | LSN | BG | FRN | CSC | TSN |
|---|---|---|---|---|---|---|---|---|
| **MALE:** | IPSI | 19.320 ± 3.761 | 15.230 ± 3.246 | 7.450 ± 2.250 | 9.763 ± 1.809 | 60.630 ± 7.904 | 47.900 ± 5.611 | |
| | CONTRA | 20.860 ± 4.509 | 11.670 ± 1.546 | 8.675 ± 2.289 | 7.979 ± 1.547 | 60.270 ± 8.431 | 46.660 ± 5.379 | |
| | P VALUE | *0.517* | *0.350* | *0.578* | *0.516* | *0.935* | *0.681* | |
| | | *Paired t-test* | *Paired t-test* | *Wilcoxon* | *Paired t-test* | *Paired t-test* | *Paired t-test* | |
| | | *t = 0.689, df = 6* | *t = 1.01, df = 6* | | *t = 0.690, df = 6* | *t = 0.086, df = 6* | *t = 0.431, df = 6* | |
| | | **HPC** | **THAL** | **LSN** | **BG** | **FRN** | **CSC** | |
| **FEMALE:** | IPSI | 21.320 ± 4.186 | 12.540 ± 3.229 | 12.030 ± 3.494 | 9.488 ± 2.934 | 63.740 ± 8.030 | 42.070 ± 10.320 | |
| | CONTRA | 20.840 ± 4.680 | 7.068 ± 2.482 | 8.889 ± 2.390 | 5.380 ± 1.862 | 45.190 ± 9.944 | 41.610 ± 10.570 | |
| | P VALUE | *0.854* | *0.163* | *0.107* | *0.023** | *0.115* | *0.813* | |
| | | *Paired t-test* | *Paired t-test* | *Paired t-test* | *Paired t-test* | *Paired t-test* | *Wilcoxon* | |
| | | *t = 0.192, df = 6* | *t = 1.590, df = 6* | *t = 1.896, df = 6* | *t = 3.020, df = 6* | *t = 1.840, df = 6* | | |
