## Supplemental Table 2 for "Seizure Circuit Activity in the Theiler’s Murine Encephalomyelitis Virus Model of Infection-induced Epilepsy Using Transient Recombination in Active Populations"

### 3 hr 4-OHT:

#### PBS VS TMEV (% area of fluorescence)

| | | HPC | THAL | LSN | BG | FRN | CSC | TSN |
|---|---|---|---|---|---|---|---|---|
| **IPSI:** | PBS | 0.869 ± 0.494 | 0.586 ± 0.208 | 1.127 ± 0.613 | 1.834 ± 0.879 | 6.376 ± 2.124 | 1.925 ± 0.565 | 5.820 ± 2.960 |
| | TMEV | 21.420 ± 2.032 | 7.666 ± 1.794 | 15.800 ± 2.262 | 3.405 ± 0.619 | 43.400 ± 5.271 | 27.07 ± 3.708 | 46.280 ± 5.201 |
| | P VALUE | *0.0001** | *0.001** | *0.0001** | *0.002** | *0.0001** | *0.0001** | *0.0001** |
| | | *Mann Whitney U* | *Mann Whitney U* | *Unpaired t-test* | *Mann Whitney U* | *Unpaired t-test* | *Unpaired t-test* | *Mann Whitney U* |
| | | | | *t = 5.590 df = 18* | | *t = 6.525, df = 18* | *t = 5.938, df = 18* | |
| **CONTRA:** | PBS | 1.927 ± 1.483 | 0.431 ± 0.182 | 0.5919 ± 0.355 | 1.050 ± 0.479 | 2.172 ± 1.124 | 1.350 ± 0.447 | |
| | TMEV | 21.830 ± 2.296 | 5.616 ± 0.890 | 15.500 ± 2.598 | 4.940 ± 1.146 | 39.200 ± 3.455 | 30.600 ± 4.636 | |
| | P VALUE | *0.0001** | *0.0001** | *0.0001** | *0.0005** | *0.0001** | *0.0001** | |
| | | *Mann Whitney U* | *Mann Whitney U* | *Mann Whitney U* | *Mann Whitney U* | *Mann Whitney U* | *Mann Whitney U* | |

| | | HPC | THAL | LSN | BG | FRN | CSC |
|---|---|---|---|---|---|---|---|
| **TMEV:** | IPSI | 21.420 ± 2.032 | 7.666 ± 1.794 | 15.800 ± 2.262 | 3.405 ± 0.619 | 43.400 ± 5.271 | 27.070 ± 3.708 |
| | CONTRA | 21.830 ± 2.296 | 5.616 ± 0.890 | 15.500 ± 2.598 | 4.940 ± 1.146 | 39.200 ± 3.455 | 30.600 ± 4.636 |
| | P VALUE | *0.862* | *0.455* | *0.579* | *0.191* | *0.372* | *0.374* |
| | | *Paired t-test* | *Wilcoxon* | *Paired t-test* | *Wilcoxon* | *Paired t-test* | *Paired t-test* |
| | | *t = 0.178, df = 12* | | *t = 0.570, df = 12* | | *t = 0.928, df = 12* | *t = 0.923, df = 12* |
| **PBS:** | IPSI | 0.688 ± 0.494 | 0.586 ± 0.208 | 1.127 ± 0.613 | 1.834 ± 0.879 | 6.376 ± 2.124 | 1.925 ± 0.5646 |
| | CONTRA | 1.927 ± 1.483 | 0.431 ± 0.182 | 0.592 ± 0.355 | 1.050 ± 0.479 | 2.172 ± 1.124 | 1.350 ± 0.4467 |
| | P VALUE | *0.688* | *0.877* | *0.375* | *0.297* | *0.297* | *0.813* |
| | | *Wilcoxon* | *Paired t-test* | *Wilcoxon* | *Wilcoxon* | *Wilcoxon* | *Wilcoxon* |
| | | | *t = 0.162, df = 6* | | | | |

#### 1.5 HR VS 3 HR 4-OHT (% area of fluorescence)

| | | HPC | THAL | LSN | BG | FRN | CSC | TSN |
|---|---|---|---|---|---|---|---|---|
| **IPSI:** | 1.5 HR | 20.320 ± 2.717 | 13.890 ± 2.231 | 9.739 ± 2.095 | 9.626 ± 1.656 | 62.190 ± 6.927 | 44.980 ± 5.699 | 55.200 ± 6.098 |
| | 3 HR | 21.420 ± 7.325 | 7.666 ± 1.794 | 15.800 ± 2.262 | 3.405 ± 0.6192 | 43.400 ± 5.271 | 27.070 ± 3.708 | 46.28 ± 5.201 |
| | P VALUE | *0.382* | *0.032** | *0.007** | *0.0007** | *0.023** | *0.007** | *0.351* |
| | | *Unpaired t-test* | *Mann Whitney U* | *Mann Whitney U* | *Unpaired t-test* | *Unpaired t-test* | *Unpaired t-test* | *Mann Whitney U* |
| | | *t = 0.382, df = 27* | | | *t = 2.985, df = 27* | *t = 2.420, df = 27* | *t = 2.938, df = 27* | |
| **CONTRA:** | 1.5 HR | 20.850 ± 3.122 | 9.370 ± 1.543 | 8.782 ± 1.590 | 6.680 ± 1.218 | 52.730 ± 6.603 | 44.140 ± 5.742 | |
| | 3 HR | 21.830 ± 2.296 | 5.616 ± 0.890 | 15.500 ± 2.598 | 4.940 ± 1.146 | 39.200 ± 3.455 | 30.600 ± 4.636 | |
| | P VALUE | *0.892* | *0.215* | *0.004** | *0.250* | *0.085* | *0.022** | |
| | | *Unpaired t-test* | *Mann Whitney U* | *Unpaired t-test* | *Mann Whitney U* | *Unpaired t-test* | *Unpaired t-test* | |
| | | *t = 0.137, df = 27* | | *t = 3.107, df = 27* | | *t = 1.786, df = 27* | *t = 2.424, df = 27* | |
